# Complete sequence of a 641-kb insertion of mitochondrial DNA in the *Arabidopsis thaliana* nuclear genome

**DOI:** 10.1101/2022.02.22.481460

**Authors:** Peter D. Fields, Gus Waneka, Matthew Naish, Michael C. Schatz, Ian R. Henderson, Daniel B. Sloan

## Abstract

Intracellular transfers of mitochondrial DNA continue to shape nuclear genomes. Chromosome 2 of the model plant *Arabidopsis thaliana* contains one of the largest known nuclear insertions of mitochondrial DNA (numts). Estimated at over 600 kb in size, this numt is larger than the entire *Arabidopsis* mitochondrial genome. The primary *Arabidopsis* nuclear reference genome contains less than half of the numt because of its structural complexity and repetitiveness. Recent datasets generated with improved long-read sequencing technologies (PacBio HiFi) provide an opportunity to finally determine the accurate sequence and structure of this numt. We performed a *de novo* assembly using sequencing data from recent initiatives to span the *Arabidopsis* centromeres, producing a gap-free sequence of the Chromosome 2 numt, which is 641-kb in length and has 99.933% nucleotide sequence identity with the actual mitochondrial genome. The numt assembly is consistent with the repetitive structure previously predicted from fiber-based fluorescent *in situ* hybridization. Nanopore sequencing data indicate that the numt has high levels of cytosine methylation, helping to explain its biased spectrum of nucleotide sequence divergence and supporting previous inferences that it is transcriptionally inactive. The original numt insertion appears to have involved multiple mitochondrial DNA copies with alternative structures that subsequently underwent an additional duplication event within the nuclear genome. This work provides insights into numt evolution, addresses one of the last unresolved regions of the *Arabidopsis* reference genome, and represents a resource for distinguishing between highly similar numt and mitochondrial sequences in studies of transcription, epigenetic modifications, and *de novo* mutations.

**Significance statement:** Nuclear genomes are riddled with insertions of mitochondrial DNA. The model plant *Arabidopsis* has one of largest of these insertions ever identified, which at over 600-kb in size represents one of the last unresolved regions in the *Arabidopsis* genome more than 20 years after the insertion was first identified. This study reports the complete sequence of this region, providing insights into the origins and subsequent evolution of the mitochondrial DNA insertion and a resource for distinguishing between the actual mitochondrial genome and this nuclear copy in functional studies.

## INTRODUCTION

Intracellular DNA transfer from mitochondrial genomes (mitogenomes) into the nucleus is pervasive and ongoing in eukaryotes (Hazkani-Covo, et al. 2010). These insertions (known as numts) are usually non-functional and subject to eventual degradation. However, they are of biological interest as a mutagenic mechanism (Turner, et al. 2003; Hazkani-Covo and Martin 2017) and the ultimate source of rare functional gene transfers from mitochondria to the nucleus (Timmis, et al. 2004). They are also of practical concern as a common cause of artifacts and misinterpretation in inferring phylogenetic relationships (Bensasson, et al. 2001), biparental inheritance of mitogenomes (Lutz-Bonengel, et al. 2021), and *de novo* mutations (Wu, et al. 2020). Most numts derive from small fragments of the mitogenome, but some can be large and structurally complex, including frequent cases where multiple discontinuous regions of mitochondrial DNA (mtDNA) fuse during integration into the nuclear genome (Portugez, et al. 2018).

The initial sequencing of Chromosome 2 in the *Arabidopsis thaliana* genome identified an extremely large numt, which was assembled to be 270 kb in length and represent approximately three-quarters of the 368 kb *Arabidopsis* mitochondrial genome (Lin, et al. 1999). However, analysis with fiber-based fluorescent *in situ* hybridization (fiber-FISH) indicated the assembly of this region was incomplete and estimated an actual size of 618 kb (± 42 kb) for the numt (Stupar, et al. 2001). This analysis suggested that large regions of repeated sequence were collapsed in the genome assembly, resulting in the erroneous exclusion of the remaining quarter of the mitogenome content that was originally inferred to be absent from the numt. Sequence comparisons between the partial numt and the *Arabidopsis* mitogenome showed high nucleotide sequence identity (99.91%), suggesting an evolutionarily recent insertion, but no evidence of selection to conserve gene function in the numt (Huang, et al. 2005).

These early analyses of the Chromosome 2 numt were hampered by multiple technical limitations. It is very difficult with conventional sequencing technologies to accurately assemble regions with long repeats that maintain high sequence identity among copies. More recent efforts to generate complete *Arabidopsis* chromosomal assemblies leveraged advances in long-read sequencing technologies (Naish, et al. 2021; Wang, et al. 2021), including PacBio HiFi, which can produce reads over 15 kb in length with >99% accuracy. These studies were successful in spanning highly repetitive centromere regions, and they both extended the coverage of the Chromosome 2 numt. However, these assemblies differed in multiple regions of the genome (Rabanal, et al. 2022), including major disagreements in the length and nucleotide sequence of this numt. The Col-CEN (Naish, et al. 2021) and Col-XJTU (Wang, et al. 2021) assemblies reported lengths of 370 kb and 641 kb, respectively, and their alignable regions differed by 109 single-nucleotide variants (SNVs), 18 indels, and one 4-bp microinversion even though they were both derived from *Arabidopsis* Col-0 ecotypes.

Another limitation in past analyses of this numt is that the original *Arabidopsis* reference mitogenome (Unseld, et al. 1997) and nuclear genome (Arabidopsis Genome Initiative 2000) derive from different ecotypes (C24 and Col-0, respectively). In addition, the original mitogenome sequence has hundreds of sequencing errors (Davila, et al. 2011; Sloan, et al. 2018). With the recent generation of accurate long-read sequencing data for the *Arabidopsis* nuclear genome (Naish, et al. 2021; Wang, et al. 2021) and a reference mitogenome for the Col-0 accession (Sloan, et al. 2018), there is a renewed opportunity to assemble and analyze this intriguing numt.

## RESULTS AND DISCUSSION

### Structure of the Arabidopsis Chromosome 2 numt

By performing a *de novo* assembly with hifiasm (Cheng, et al. 2021) of PacBio HiFi reads generated as part of the recent Col-CEN effort to span the centromeres in the *Arabidopsis* genome (Naish, et al. 2021), we produced a gap-free contig that covered the entire numt insertion in Chromosome 2 (**Figure 1**). The large numt was embedded within a 12.6-Mb contig and was consistent in both size (641-kb) and structure with the recent Col-XJTU genome assembly (Wang, et al. 2021), but it differed considerably in nucleotide sequence (see below). Our assembly also matched the repeat structure previously inferred from fiber-FISH and fell within the estimated size range of 618 ±42 kb from that analysis (Stupar, et al. 2001).

**Figure 1.**
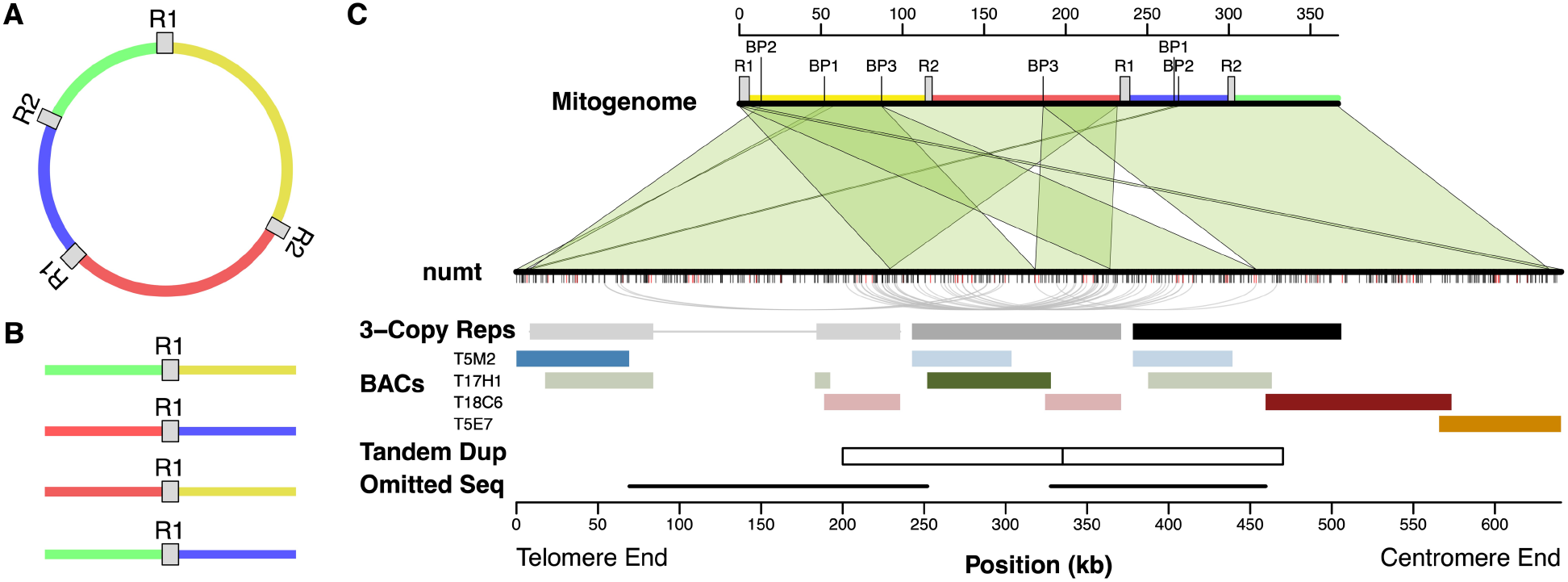
Structure of the *Arabidopsis* Chromosome 2 numt. (A) A simplified circular representation of the *Arabidopsis* mitogenome. The sequence from the C24 ecotype was used for structural comparisons with the numt because the Col-0 mitogenome contains rearrangements associated with recombination at small repeats (see main text). This conformation of the C24 mitogenome corresponds to the previously described D’–A’–C–B structure (Stupar, et al. 2001). R1 and R2 indicate the two large pairs of repeats. Intervening single-copy regions are in different colors, also indicated on the mitogenome map in panel C. (B) Recombination between a pair of repeats in the mitogenome produces four possible alternative combinations of flanking sequences (as shown for R1) which are thought to be present at near equal frequencies in tissue samples. The first three of these conformations are all found within the numt. (C) Structural comparison of the numt and mitogenome. The mitogenome sequence (top) is annotated with the large repeat sequences (R1 and R2) and pairs of breakpoints (BP1, BP2, and BP3) associated with chimeric fusions in the numt that are possibly the result of non-homologous end-joining. Green shaded regions show blocks of syntenic sequence conserved in the numt (bottom). Tick marks below the numt show SNVs (black) and indel/structural variants (red) relative to the Col-0 mitogenome sequence. Some large sections of the mitogenome appear three times in the numt (indicated in shades of gray to black in the 3-Copy Reps row). The curved gray lines connect pairs of variants where two copies share an allele that differs from the mitogenome and the other repeat copy. The colored blocks show locations of four bacterial artificial chromosomes (BACs) originally used to assemble this genome region. The darker block for each BAC indicates the actual location of that BAC within the numt. The blocks in fainter colors represent repeated sequences similar to the BAC. The adjacent white boxes (Tandem Dup) represent the resulting copies from a putative 135-kb tandem duplication that occurred within the nuclear genome after the numt had already begun to diverge in sequence. The repetitive structure of the numt led to the T17H1 BAC being incorrectly overlapped with the T5M2 and T18C6 BACs in the original *Arabidopsis* genome assembly, resulting in the exclusion of two large regions of intervening sequences (indicated by the black lines in the Omitted Seq row).

The assembled numt is considerably larger than the reference *A. thaliana* Col-0 mitogenome because of extensive sequence duplication, including large tandem repeats. The earlier fiber-FISH study (Stupar, et al. 2001) concluded that repeat-mediated overlap between bacterial artificial chromosomes (BACs) used in the original nuclear genome assembly led to the exclusion of a single large internal region. However, by obtaining the entire numt sequence, we found that the T17H1 BAC does not represent the repeat on the centromere-end of the numt as previously inferred. Instead, this BAC derives from the middle of three repeat copies in the numt, meaning that two flanking regions on either side of the T17H1 BAC were omitted from the original assembly (**Figure 1c**).

The numt also exhibits multiple structural differences relative to the mitogenome, including rearrangements arising from recombination between two different pairs of small repeats, which are known as the C and Q repeats and are 457 and 206 bp in length, respectively (Davila, et al. 2011) (**Figure S1**). Even though the *A. thaliana* nuclear genome sequence derives from the Col-0 ecotype, the conformations associated with these repeat pairs match the *A. thaliana* C24 mitogenome (Unseld, et al. 1997). Therefore, the repeat-mediated recombination events that distinguish the Col-0 and C24 mitogenomes likely occurred in the Col-0 mitogenome after the numt insertion, consistent with the relatively rapid accumulation of these rearrangements in the divergence of mitogenome structures among *Arabidopsis* ecotypes (Arrieta-Montiel, et al. 2009). However, it is also possible that occasional outcrossing within this largely selfing species (Platt, et al. 2010) has led to discordance between the genealogies of the numt and the mitogenome, such that the Col-0 numt is more closely related to the C24 mitogenome than the Col-0 mitogenome.

The *Arabidopsis* mitogenome also contains two pairs of large repeats (6.0 and 4.2 kb in size). In plant mitogenomes, repeats of this size undergo near-constant recombination such that they are present in multiple alternative structures, even within tissue samples (Gualberto and Newton 2017). Three of the four possible alternative conformations associated with the “Repeat 1” pair are found in the numt, meaning that the same flanking sequence can have two different connections on the other side of the repeat (**Figure 1**). We infer that these alternative structures result from the direct transfer of multiple copies from the mitogenome. Although it is possible that rearrangements generated them within the nucleus after insertion, the fact that the alternative structures already exist at high frequencies within the mitochondria makes direct transfer a much more likely explanation. Therefore, some of the repetitiveness of this complex numt appears to result from the original transfer. Mitogenomes are known to exist in complex structures, including multimeric forms (Bendich 1993), so it is possible that a single transferred molecule could have contained multiple copies of some regions, including these alternative structures. However, complex numts commonly arise via fusion of multiple DNA fragments (Portugez, et al. 2018), so it is also possible that the alternative structures were present in distinct DNA fragments that fused at the time of insertion.

Although most of the numt shows conserved synteny with the reference mitogenome or can be explained by repeat-mediated recombination events (see above), there are also structural rearrangements with breakpoints that appear to result from non-homologous end joining (NHEJ). The first 8 kb of sequence at the telomere-end of the numt consists of two fragments from disparate parts of the mitogenome that appear to result from fusion events (BP1 and BP2 in **Figure 1c**). In addition, there is an internal breakpoint in the numt that is not associated with repeat sequences in the mitogenome (BP3 in **Figure 1c**). This novel fusion is duplicated within the numt as part of a large tandem repeat structure. As discussed below, the patterns of sequence divergence among these repeats provide insight into the further expansion of the numt after its original insertion.

### History of nucleotide sequence divergence in the Arabidopsis Chromosome 2 numt

Even though the structure and length of our numt assembly generally match the corresponding regions in the recent Col-XJTU assembly, the two assemblies differ substantially in sequence. Most notably, the Col-XJTU numt sequence has 260 SNVs relative to our assembly (**Table S1**). In every one of these cases, the Col-XJTU variant matches the Col-0 mitogenome even in the large regions of the assembly where BACs provide independent validation of our basecalls (**Figure 1**). Therefore, large portions of the Col-XJTU numt assembly appear to have been “overwritten” by the more-abundant reads derived from the highly similar mitogenome sequence. To further investigate the sequence discrepancies with the Col-XJTU assembly, we performed a *de novo* assembly of the Col-XJTU HiFi reads, which generated a near-identical sequence (differing by only 5 SNVs) to our *de novo* assembly of the Col-CEN HiFi reads. Read mapping indicated that these SNVs reflect true differences between the samples used for Col-CEN and Col-XJTU projects (**Table S2**). Accordingly, the Col-XJTU project identified >1000 sequence variants and/or errors genome-wide (Wang, et al. 2021), suggesting some divergence among the sequenced Col-0 lines.

By comparing the numt to the reference Col-0 mitogenome, we found that they were 99.933% identical in nucleotide sequence (after excluding indels, multinucleotide variants, and short unalignable sequences adjacent to indel regions). This level of sequence identity is even higher than a previously reported value of 99.91% (Huang, et al. 2005), which is not surprising because that study was based on only a portion of the numt and a C24 mitogenome reference that was since found to contain numerous sequencing errors. The SNVs that distinguish the numt and the mitogenome are dominated by transitions with GC base-pairs in the mitogenome and AT base-pairs in the numt (**Tables 1 and S3**). This signature likely reflects the much higher rate of mutation in the nuclear genome than the mitogenome (Wolfe, et al. 1987; Drouin, et al. 2008) and the biased mutation spectrum in the nucleus (Ossowski, et al. 2010; Weng, et al. 2019). SNV transitions showed a bias of 6.7 to 1 towards AT base-pairs in the numt. This bias is approximately twice as strong as previously reported (Huang, et al. 2005), indicating that our improved numt assembly and a higher quality mitogenome reference have substantially reduced noise. The sequence divergence between the numt and the mitogenome also showed evidence of a deletion bias in the nuclear genome (Weng, et al. 2019), as more than two-thirds of the indels that distinguished the two genomes had the shorter allele in the numt (**Table 1**).

**Table 1.** Sequence variants distinguishing the *Arabidopsis* Chromosome 2 numt from the Col-0 reference mitogenome sequence

| Variant | Count |
| --- | --- |
| **Total SNVs (Mitogenome<>numt)** | **425** |
| **Total Transitions** | **270** |
| GC<>AT | 235 |
| AT<>GC | 35 |
| **Total Transversions** | **155** |
| GC<>TA | 58 |
| AT<>CG | 30 |
| GC<>CG | 42 |
| AT<>TA | 25 |
| **Total Indels** | **44** |
| numt shorter | 30 |
| numt longer | 14 |

The C➔T transitions that dominate the numt mutation spectrum are a hallmark of the abundant 5-methylcytosine (5mC) modifications at CpG and CHG sites in plant nuclear genomes (Vanyushin and Ashapkin 2011; Weng, et al. 2019; Naish, et al. 2021; Monroe, et al. 2022). We found that 88 of the 235 C➔T observed SNVs occur at CpG sites, and an additional 87 occur at CHG sites. This total of 74.5% (175 of 235) represents a highly significant enrichment relative to the 33.3% of all cytosines in the mitogenome that are found in a CpG or CHG context (χ^2^ = 178.9; *p* < 0.0001), supporting the expected role of 5mC modifications in numt sequence divergence. Furthermore, using previously generated nanopore sequencing data (Naish, et al. 2021), we found high levels of 5mC modifications across the full-length of the numt, consistent with observations for pericentromeric regions in the rest of the *Arabidopsis* genome (**Figure 2**). This high level of methylation supports previous conclusions that the numt is likely to be transcriptionally inactive (Huang, et al. 2005; Adamo, et al. 2008).

**Figure 2.**
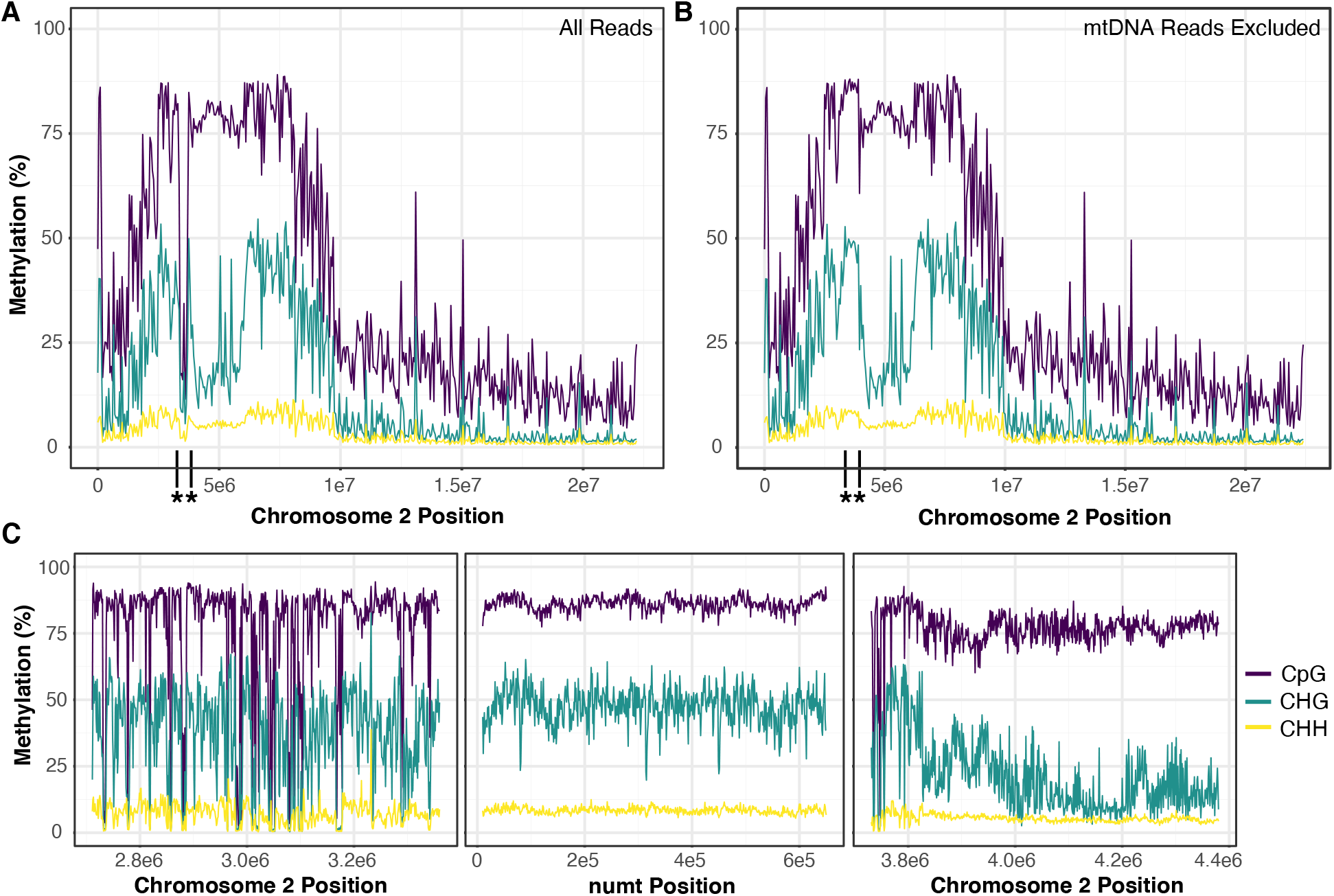
Nanopore-derived estimates of methylation percentage across Chromosome 2 of the Col-CEN assembly (after updating it to include the full numt) in CpG (purple), CHG (teal) and CHH (yellow) contexts. (A) Methylation profile including all reads (>30 kb) averaged over 50-kb windows. The boundaries of the numt region are indicated with asterisks and vertical black lines on the x-axis. (B) The same profile after excluding mitogenome-derived reads based on SNVs that distinguish the numt and mitogenome, which greatly increases the estimated methylation levels in the numt because of the lack of methylation in the actual mitogenome. (C) Methylation profile of 650 kb on the telomere side of the numt (left) across the numt (middle) and 650 kb on the centromere side of the numt (right) averaged over 1-kb windows.

The repetitive structure in the numt raises the possibility of a large duplication that occurred during the initial insertion event or one that occurred within the nuclear genome post-insertion. We reasoned that patterns of sequence divergence could differentiate between these alternative models (Hazkani-Covo, et al. 2003). If duplicates were generated at the time of insertion, all copies will have started diverging simultaneously and form a “star phylogeny”. In contrast, later duplications within the nucleus after sequence divergence had already begun would lead to descendent copies sharing derived variants with each other. Therefore, we compared sequence divergence among the large repeat regions present in three copies in the numt (**Figure 1c**) and the homologous mitogenome sequence. We found a higher average pairwise divergence between repeats within the numt (0.095%) than between those sequences and the reference mitogenome (0.065%). Again, this is consistent with a higher mutation rate in the nucleus than in the mitogenome. We identified 34 variants for which one of the three copies in the numt matched the mitogenome reference and the other two shared an alternative allele (**Figure 1c, Table S4**). Given the extremely low rate of sequence divergence between repeats, these patterns of shared alleles are highly unlikely to arise by independent mutations (i.e., homoplasy). Instead, they suggest a duplication after nucleotide sequence divergence had already started to occur following the initial numt insertion.

Most of these shared variants occurred in a consistent fashion, supporting tandem duplication of a 135-kb sequence, with a central breakpoint at ~335 kb from the telomere end of the repeat (**Figure 1c**). However, a cluster of four variants shows a conflicting pattern, linking the internal duplicated region with repeated sequence content at the far telomere end of the numt (**Figure 1c**). These pairings are more difficult to interpret but could reflect a history of localized gene conversion after repeat copies began to diverge. Comparing the divergence of this numt sequence among closely related *A. thaliana* ecotypes may help further tease apart the effects and timing of gene conversion and duplication events.

In summary, the accuracy of PacBio HiFi technology can resolve extremely complex genome structures consisting of long repeats that share highly similar (but non-identical) sequences. *Arabidopsis* is the pre-eminent model system in plant genetics, so obtaining complete and accurate genomic resources is of utmost importance. The original *Arabidopsis* genome assembly (conducted more than two decades ago; Arabidopsis Genome Initiative 2000) and recent efforts to close the remaining centromere-based gaps (Naish, et al. 2021; Wang, et al. 2021) represent major landmarks in that process. The resulting PacBio HiFi sequencing data have allowed us to address one of the last remaining unresolved regions in the genome assembly. To our knowledge, this represents the largest numt ever sequenced. Large numt tandem arrays have recently been identified in humans and can reach similar sizes (Lutz-Bonengel, et al. 2021), but they have yet to be sequenced. Smaller numt fragments have also undergone massive proliferation into large tandem arrays in legumes (Choi, et al. 2022). Insertions of near-complete genomes of plastids and other bacterial endosymbionts have also been observed (Huang, et al. 2005; Dunning Hotopp, et al. 2007). Therefore, these large insertions are likely common elements of eukaryotic genomes that are frequently overlooked because of challenges associated with assembling regions with such high similarity to organelle/endosymbiont genomes.

Numts are a source of fascination because of their biological importance but also frustration as a source of artifacts in genetic studies. In addition to providing insights into the origins and evolution of this extremely large and complex numt, a complete sequence of this region is of practical value for distinguishing between the numt and true mtDNA in studies investigating molecular processes such as *de novo* mutation, transcriptional activity, and epigenetic modifications. The similarity of the numt and mitogenome will still pose challenges (especially for short-read sequencing technologies) because stretches of thousands of base-pairs remain 100% identical between the numt and the mitogenome, but the set of reliable variants (**Figure 1, Table S3**) provides a foothold for distinguishing molecular processes associated with these highly similar sequences.

## METHODS

### De novo genome assembly

To generate a *de novo* assembly of the numt region, we used the full set of PacBio HiFi reads (circular consensus sequences) from Naish et al. 2021, which were accessed via the European Nucleotide Archive (accession number PRJEB46164) on Nov. 18, 2021. We used the hifiasm v. 0.15.1-r334 assembler (Cheng, et al. 2021), which was developed for the specific purpose of assembling long, highly accurate reads such as those from PacBio HiFi sequencing. Because the focal genotype is highly inbred, we included the ‘-l0’ flag as part of the assembler configuration, thereby disabling automatic duplication purging. The resultant assembly graph was converted to a set of contigs in a multi-fasta format using AWK (Aho et al. 1988) as described at https://github.com/chhylp123/hifiasm. To identify the numt region in the resulting contigs we used a local BLAST database (Altschul et al. 1990) and a query composed of the previous, partial assembly of the *A. thaliana* numt sequence. We later repeated these assembly methods with an independent PacBio HiFi dataset (Wang, et al. 2021), accessed via the Genome Warehouse in the National Genomics Data Center, Beijing Institute of Genomics, Chinese Academy of Science / China National Center for Bioinformation (BioProject PRJCA005809) on Nov. 28, 2021. The structural accuracy of the assembly was validated using multiple orthogonal approaches, including alignment consistency of published Illumina, PacBio HiFi, and nanopore reads mapped to the assembled sequences (Naish, et al. 2021; Wang, et al. 2021), consistency with the published BAC sequences (Lin, et al. 1999), consistency with published fiber-FISH results (Stupar, et al. 2001), and consistency with published BioNano optical mapping data (Naish, et al. 2021).

### Comparative sequence analysis

EMBOSS Stretcher (https://www.ebi.ac.uk/Tools/psa/emboss_stretcher/) was used to generate global pairwise alignments between different assemblies of the numt region. In addition, this aligner was used to compare our assembly to a manually generated rearrangement of the Col-0 mitogenome (GenBank accession NC_037304.1), for which homologous regions of the mitogenome were concatenated to match the synteny of the numt. Multiple sequence alignments of the large repeats in the numt (**Figure 1c**) and homologous mitogenome sequence were generated with MAFFT v7.453 under default parameters. Variants in aligned sequences were identified and quantified with custom Perl scripts. Sequence variants and structural comparisons between the numt, mitogenome, and BACs from the original *Arabidopsis* genome project were visualized with a custom script run in R v4.0.5.

We assessed the quality of basecalls in the *de novo* numt assembly with local BLAST alignments of the assembly against the numt derived BACs from the original *Arabidopsis* genome assembly and identified 7 SNVs distinguishing the *de novo* assembly and the BACs (**Table S5**). To validate these 7 SNVs, we aligned the HiFi reads to the *de novo* numt assembly using minimap2 v. 2.22 (Li 2018) and manually inspected the alignments using IGV (Thorvaldsdóttir, et al. 2013). For all 7 SNVs, the HiFi reads unanimously supported the allele in the *de novo* numt assembly. We also used the mapped HiFi reads to manually confirm support for 5 observed SNVs that distinguished our *de novo* assemblies of the Col-CEN and Col-XJTU HiFi reads (**Table S2**).

### Cytosine methylation analysis

Previously published nanopore reads (Naish, et al. 2021) were filtered for length (>30kb) using Flitlong (--min_mean_q 95, --min_length 30000; https://github.com/rrwick/Filtlong) and aligned to our *de novo* Col-CEN numt assembly and the reference Col-0 mitogenome using Winnowmap v1.11, -ax map-ont) (Jain, et al. 2020). Alignments were filtered for those containing the numt allele at each SNV position (**Table S3**) using SplitSNP (https://github.com/astatham/splitSNP). Bam files were merged using Samtools v1.9 and read IDs were extracted and filtered to retain only duplicate IDs (>2). The resulting readset was used for methylation calling against the numt assembly with Deepsignal-plant v0.14 (Ni, et al. 2021). Whole-chromosome methylation analysis was performed with the full 30-kb dataset and with the dataset generated by removing reads containing mitogenome alleles.

## Data and code availability

All scripts are available via https://github.com/dbsloan/arabidopsis_numt. Alignments and numt sequences are available via https://zenodo.org/record/6168939.

## ACKNOWLEDGEMENTS

This work was supported by grants from the National Institutes of Health (R01 GM118046) to D.B.S., the Human Frontier Science Program (RGP0025/2021) to M.C.S. and I.R.H., and the Biotechnology and Biological Sciences Research Council (BB/V003984/1) to I.R.H.

## SUPPLEMENTAL MATERIAL

**Figure S1.**
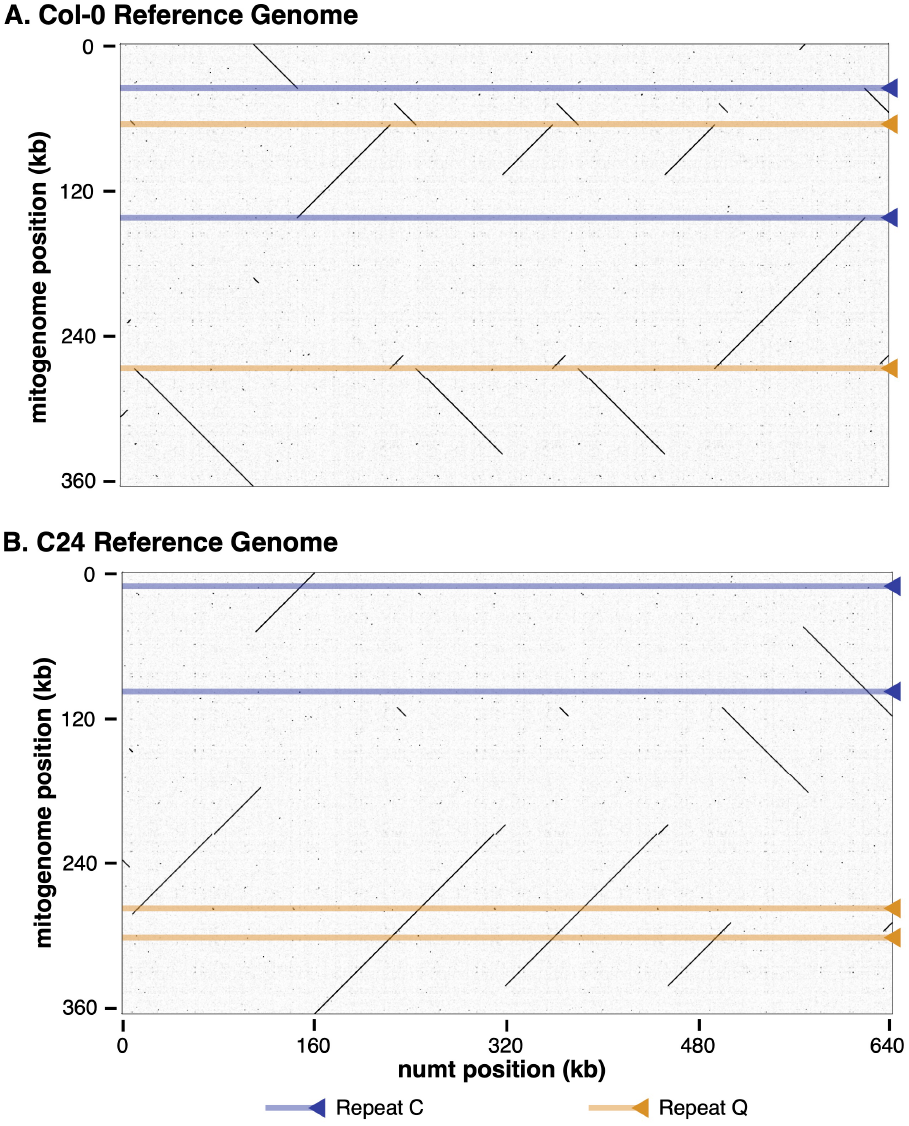
Dot plots comparing structure of the *A. thaliana* Chromosome 2 numt to the published reference mitogenomes for (A) *A. thaliana* Col-0 (NC_037304.1) and (B) *A. thaliana* C24 (Y08501.2). Black diagonal lines indicate regions of conserved synteny between the numt and the corresponding mitogenome. The positions of the C and Q repeats in the mitogenome are highlighted in blue and orange, respectively. Note that these two repeats are associated with breaks in conserved synteny with the Col-0 mitogenome due to repeat-mediated recombination but not with the C24 mitogenome. Dot plots were generated with gepard v2.1.0 (Krumsiek, et al. 2007).

**Table S1.** SNVs that distinguish the numt in our *de novo* assembly of Col-CEN HiFi reads from the corresponding sequence in the published Col-XJTU assembly. Position numbering is relative to the telomere end of the numt in our *de novo* assembly.

**Table S2.** Variants that distinguish the numt in our *de novo* assembly of Col-CEN HiFi reads from our *de novo* assembly of the Col-XJTU HiFi reads. Position numbering is relative to the telomere end of the numt in the *de novo* Col-CEN assembly.

**Table S3.**Variants that distinguish the numt in our *de novo* assembly of Col-CEN HiFi reads from the reference Col-0 mitogenome (NC_037304.1). Position numbering is relative to the telomere end of the numt.

**Table S4.** Pairs of sites in the 3-copy repeats that share a different allele than the mitogenome and the other repeat copy. Position numbering is relative to the telomere end of the numt.

**Table S5.** Variants that distinguish the numt in our *de novo* assembly of Col-CEN HiFi reads from the sequenced BACs in the original *Arabidopsis* genome project. Position numbering is relative to the telomere end of the numt.

